# Regional neutrality evolves through local adaptive niche evolution

**DOI:** 10.1101/326348

**Authors:** Mathew A. Leibold, Mark C. Urban, Luc De Meester, Christopher A. Klausmeier, Joost Vanoverbeke

## Abstract

Joost Vanoverbeke, Department of Biology, Katholieke Universiteit Leuven, Leuven, Belgium. Abstract:
Biodiversity in natural systems can be maintained either because niche differentiation among competitors facilitates stable coexistence or because equal fitness among neutral species allows for their long-term co-occurrence despite a slow drift toward extinction. Whereas the relative importance of these two ecological mechanisms has been well-studied in the absence of evolution, the role of local adaptive evolution in maintaining biological diversity through these processes is less clear. Here we study the contribution of local adaptive evolution to coexistence in a landscape of interconnected patches subject to disturbance. Under these conditions, early colonists to empty patches following disturbance can often adapt to novel local conditions sufficiently fast to prevent successful colonization by other pre-adapted species. Over the long term, the iteration of these local-scale priority effects results in niche convergence of species at the regional scale even though species tend to monopolize local patches. Thus, the dynamics evolve from stable coexistence through niche differentiation to neutral co-occurrence at the landscape level while still maintaining strong local niche differentiation. Our model results show that neutrality can emerge at the regional scale from local, niche-based adaptive evolution, potentially resolving why ecologists often observe neutral distribution patterns at the landscape level despite strong niche divergence among local communities. Our results also demonstrate how local adaptive evolution can shape cryptic eco-evolutionary dynamics and thus alter the regional mechanisms that determine biological diversity and resistance to disturbance.

## Significance statement

Coexistence of species depends on two very general mechanisms. In one, species differentiate in their niches and coexist by negative frequency dependence, in the other they have similar niches and cooccur for long periods of time due to stochastic processes. These explanations ignore the role of local adaptive evolution. We model how local evolutionary priority effects (where species that recolonize patches can adapt sufficiently well to resist being displaced by later colonists) affect coexistence in a landscape and find that evolution often leads to a situation where local patches are strongly dominated by a single species even though all species can be found in any habitat type. This unsuspected result may explain why coexistence of species is often scale dependent.

## Introduction

Biologists have long sought to understand what maintains the vast diversity of life on Earth (Hutchinson 1959). Two major hypotheses have emerged: either species respond to environmental variation through distinct niche differences or they are equivalent in their niches and drift via random processes. The maintenance of biological diversity among competing species is consequently increasingly understood as a tension between the stabilizing properties of niche differences versus fitness equality without niche differentiation, which results in neutral drift (Chesson 2000, Leibold and McPeek 2006, Adler et al. 2007). Niche-based mechanisms assume that competing species differ in their niches such that each species affects its own reproductive success more than that of other species and can increase when rare in the overall environment. This dynamic favors the coexistence of species with different niches. The neutral mechanism allows for long-term co-occurrence of species with equivalent fitness across environments despite the lack of stabilizing ecological feedbacks; eventually though, through random drift, one species will dominate and the other species will slowly become extinct. Is the diversity that we observe in nature determined by niche differentiation or similarity? The answer to this question has profound implications for how we understand the distribution of coexisting species and how to best maintain biodiversity in the face of natural or anthropogenic disturbances.

Previous investigations of this question have so far assumed that stable niche-based coexistence and drift can both operate and that either can predominate depending on landscape features and average species traits (e.g. Gravel 2006, Leibold and McPeek 2006, HilleRisLambers et al. 2012). However, these approaches have generally ignored the role of local adaptive evolution.

While adaptive evolution has often been seen as a slow process relative to ecological dynamics, empirical work increasingly shows that local adaptive evolution can occur rapidly (Tessier et al. 1992, Hendry & Kinnison, 2001; Ellner et al. 2011), and that it can occur at relatively fine spatial scales (Richardson et al 2014; Reznick 1998), thus intersecting the same temporal and spatial scales over which stabilizing and equalizing ecological mechanisms act. Furthermore, the strength of evolutionary change on population dynamics and community composition can rival or even exceed those due to purely ecological change such as abiotic effects or disturbances (Ellner et al. 2011, Pantel et al. 2015). Evolution can thus act as an underappreciated effect that can affect ecological patterns in nature (Yoshida, 2007; Urban, 2013; Hiltunen 2014).

Past theoretical work in this area suggests that, depending on assumptions, the effects of local adaptation can either cause competing species to diverge (Schluter & McPhail 1993) or converge (Riley 1963, Aarssen 1986, Hubbell 2006, Scheffer and van Nes 2006, terHorst et al. 2010) in niche traits, facilitating niche partitioning or neutral co-occurrence of species, respectively. This research, however, neglects the regional scale and the process by which communities assemble through repeated colonization and competition. Taking this more regional perspective, local adaptive evolution can generate evolution-mediated priority effects wherein early colonizers adapt to local environmental conditions, monopolize local resources, and prevent invasion by, and subsequent local coexistence with, later colonizers (De Meester et al. 2016). This could result in species having narrow, dedicated niches in each local environment (local adaptation), but broad overlapping niches at the regional scale (neutrality). This outcome likely depends on the species’ relative dispersal rates within a landscape of patches, or metacommunity (sensu Leibold et al. 2004), as dispersal links local eco-evolutionary dynamics with regional processes involving disturbance, and habitat variability (Urban et al. 2008, Loeuille and Leibold 2008, Vanoverbeke et al. 2016).

Here we study the effects of eco-evolutionary feedbacks on niche versus neutral processes at local and regional scales. We first combine an analytically tractable, albeit highly simplistic, deterministic patch dynamics model of community assembly with a simple model of adaptive dynamic evolution. This model highlights several unique features of the role of adaptive evolution in metacommunities. We then study a more realistic spatially explicit individual-based model where we explore how robust this solution is to dispersal rate, disturbance frequency, and reproductive system (sexual vs asexual) as well as stochasticity due to demographic and dispersal effects in finite communities. We also apply common ecological tools to our simulations and find that local niche divergence can lead to regional neutrality while simultaneously maintaining relatively strong local niche partitioning, thus providing a more integrated view of the mechanisms underlying biodiversity maintenance and possibly explaining why many large scale empirical studies find neutral-like patterns of species distributions in nature even in systems that are thought to have high niche differentiation.

### Patch occupancy model of community assembly

As a first step toward incorporating local adaptation, we develop a minimalist patch dynamics model that describes community assembly and adaptation as a series of transitions among occupation and adaptive states (Law and Morton 1993). In this model, alternate species of potential colonizers undergo an iterated process of colonization and extinction depending on disturbance frequency, local environmental conditions, resident species, and colonist traits.

Conventional models of community assembly assume that species have fixed ecological niche attributes and that this process thus can be understood by describing the rules that govern the set of possible successful colonizations and their consequent effects on extinction of other species in a deterministic way (e.g. Law and Morton 1993, 1996, Law and Leibold 2005). These rules can be described by a community assembly graph as shown in Figure 1a where they are applied to a simple model of community assembly for two strongly competing species in two alternate habitat types in a scenario where each species is competitively superior in different habitats (a “harlequin landscape”, *sensu* Horn and MacArthur 1972). Local adaptation can be included by incorporating additional rules that describe species transition from maladapted to adapted states and how these states alter further transitions among species (Figure 1b).

**Figure 1:**
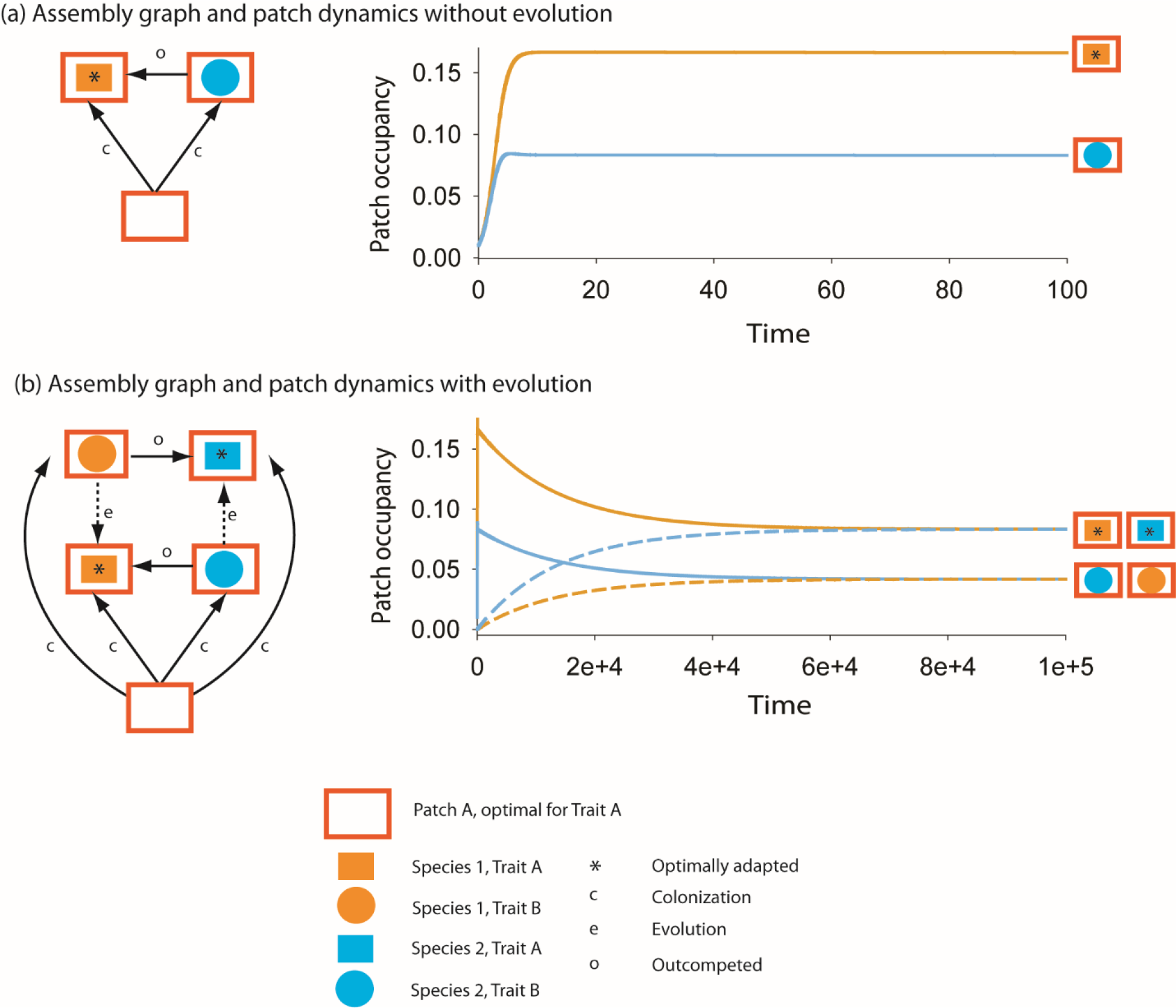
Evolution alters community assembly by creating neutrality between species at the metacommunity level. We present assembly graphs (left) and results of patch occupancy models (right) for patch A. The structure and dynamics for patch B (not shown) mirror those for patch A. In (a) we assume no evolution and in (b) we assume that both species can adapt to the other species’ niche. In the assembly graphs on the left, colors indicate species identity (orange for species 1 and blue for species 2). Shapes indicate patch type and trait value. For instance, trait A (orange filled rectangle) is optimal in patch A (orange open rectangle). Asterisks denote occupancy stages that are not invasible and thus end-points for community assembly because they are occupied by optimally adapted trait types, as denoted by the matching of rectangle traits to rectangle environments and an asterisk. Not shown are local extinctions due to disturbances that revert any occupied patch back to empty. Evolution in (b) considerably expands the number of possible species-trait-patch combinations. Arrows indicated by a c denote transitions that occur due to colonization, those indicated by an o denote transitions that occur when a colonist species outcompetes and excludes a resident one and dotted lines indicated by e’s indicate evolution into a different trait type by a resident species. On the right, we show the respective patch dynamics of each assembly graph for patch A. Without evolution (a), type A patches are dominated by pre-adapted species 1, but maladapted species 2 also occurs in recently disturbed patches as recent colonists from other patches until species 1 arrives. With evolution, the initial dynamics are almost identical to what happens without evolution for the first 100 time steps but then slowly change as shown in (b) (note the change in time scale) and eventually change so that both species occur in patch A in their adapted state and at equal frequencies. Maladaptive phenotypes of both species also occur at lower densities through colonization from patch B as shown by the dotted lines.

**Table 1:** Parameters of the patch occupancy model.

| **parameter** | **Value** | **Description** |
| --- | --- | --- |
| $h_A, h_B$ | 0.5 | |
| $c_{ad}, c_{mal}$ | 2 | |
| $m$ | 1 | |
| $\mu$ | $1e^{-4}$ | |

**Table 2:**
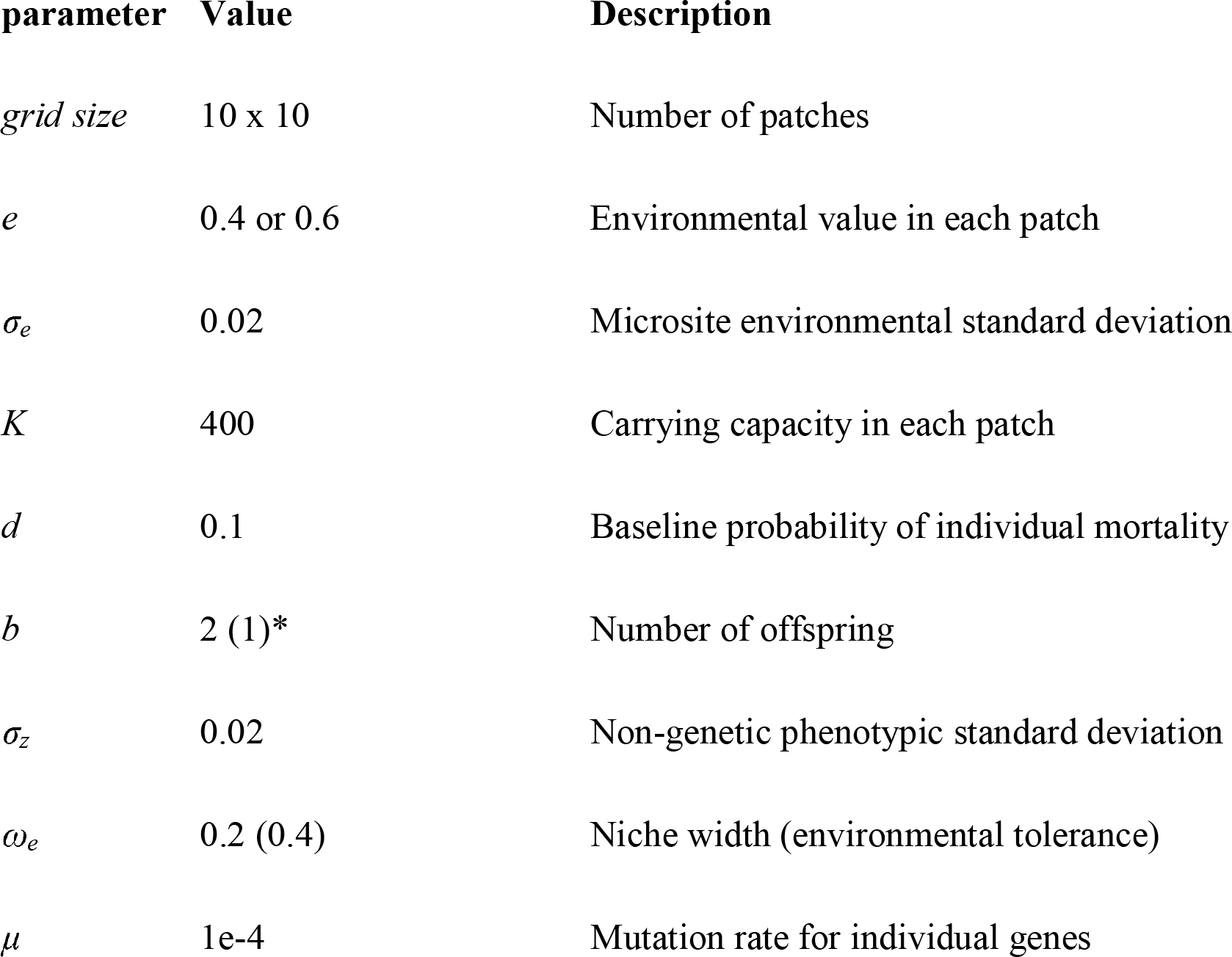
Parameters of the IBM.

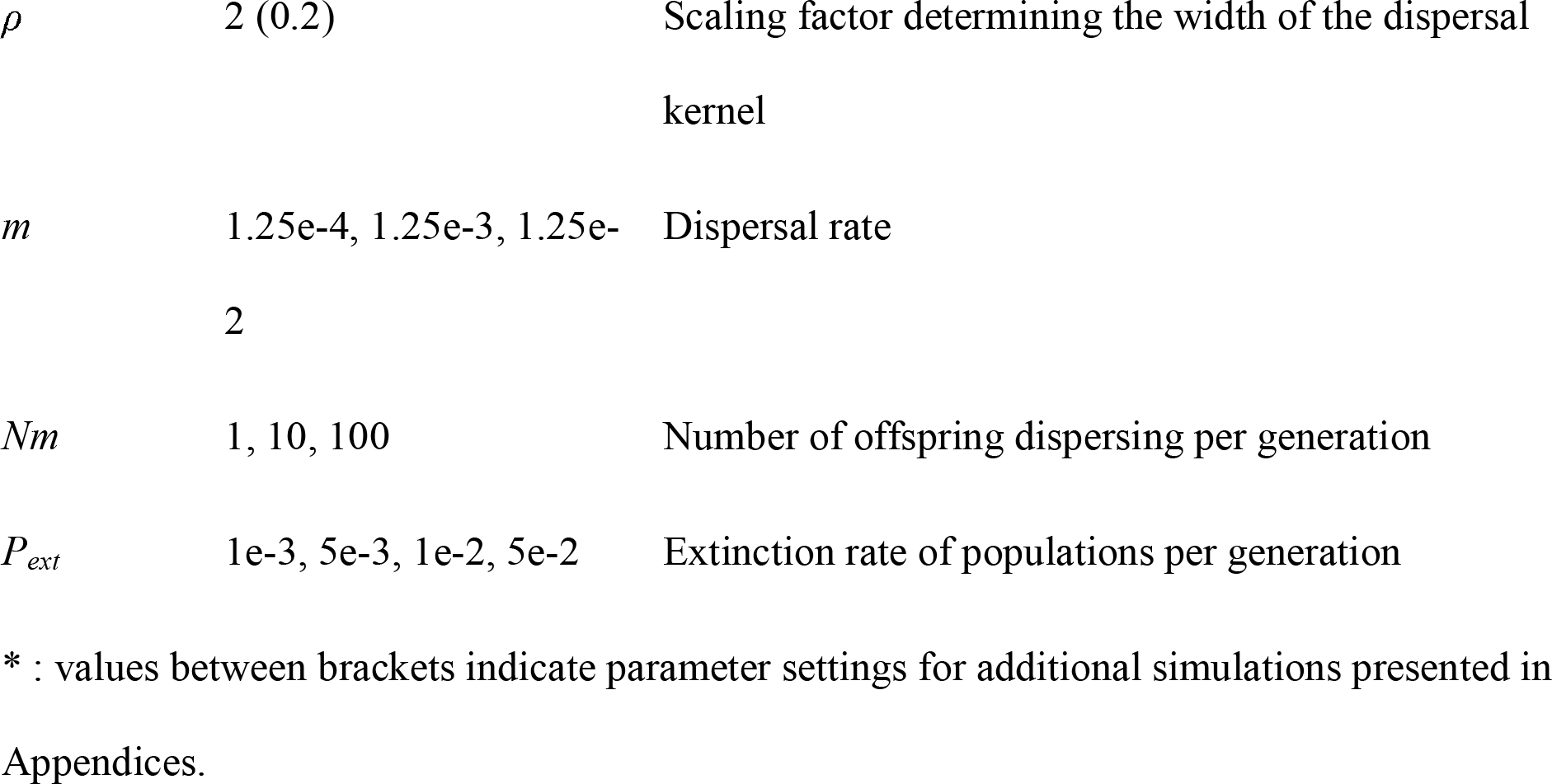

Using these rules, we develop a model of two competing metapopulations. Each population can be adapted to one or the other habitat type. Local adaptation occurs as a state transition. We allow for evolution-mediated priority effects by disallowing local coexistence and assuming that the locally adapted resident (regardless of species identity) cannot be invaded (Fig. 1b). To minimize the possibility that regional neutrality is important we initiate our simulations with two species each of which is adapted to and completely dominant in each of two alternate patch types and ask how subsequent evolution may alter their niche relations and spatial distributions. We find that in the short term, both species continue to strongly segregate by patch type (Figure 2a) but that little by little, each species is increasingly found in the alternate patch type (Figure 2b). Eventually, both species are equally distributed in both patch types (Figure 2b) even though (by assumption) they never coexist in any local patch.

**Figure 2:**
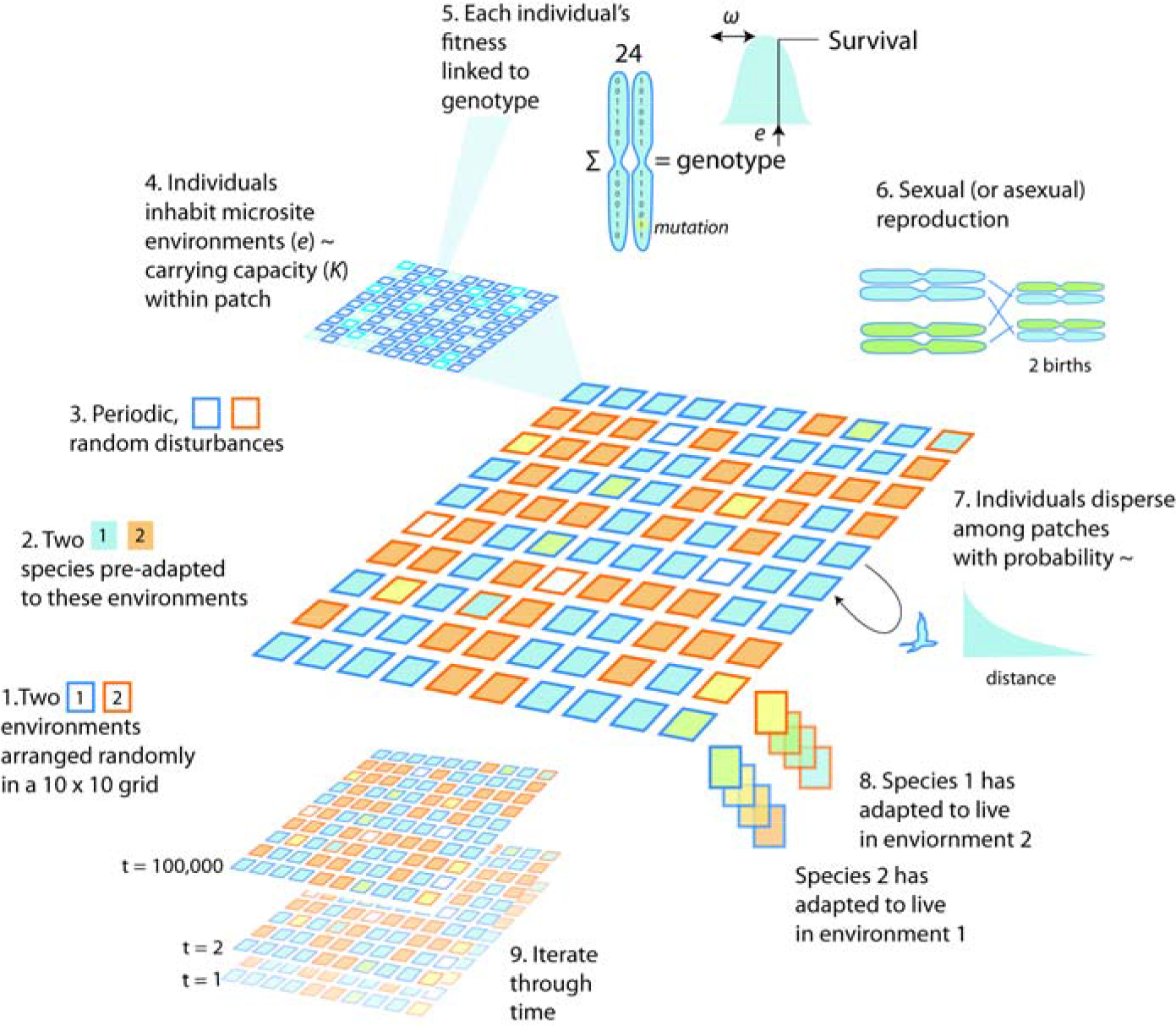
Schematic illustration of the simulation model. Each simulation starts with species that are optimally adapted to alternate habitat types in patches arrayed in a grid landscape. Each patch has a carrying capacity and populations are maintained through either sexual or asexual reproduction. Random extinctions of the entire population of individuals in patches creates empty patches. Mutations generate ecotypic variation among individuals that is subject to local selection depending on the patch type. Dispersal leads to recolonization of empty patches and gene flow among patches with existing populations. This process is iterated through time and allows each species to become more adapted to alternate patch types than it was at the beginning of the simulation. We find that eventually both species can become adapted to both patch types at rates and frequencies that depend on model parameters.

This model serves to illustrate some of the key elements of the process that could generate what we define as regional neutrality. These elements include 1) the evolutionary convergence of similar bimodal niche trait distributions at the regional scale, 2) convergence of species distribution among patch types toward the frequency distribution of patch type, and 3) erosion of correlations between species distribution and habitat type after the convergence of trait distributions.

### Spatially explicit IBM of community assembly

To study how regional neutrality could evolve under more realistic assumptions, we used individual-based simulations to model a similar scenario on a spatially explicit 10 by 10 grid of patches with two patch types and two species (Figure 2). Our simulations allowed us to manipulate dispersal rate, disturbance rate, strength of selection, and reproduction type (sexual vs asexual). Evolution was modeled by assuming that competitive dominance was determined by a single multilocus trait with different optimal values for each alternative habitat type. Genetic variance in this trait was modeled by assuming zero initial standing genetic variation and allowing for random mutations at each locus to generate and maintain genetic variation in subsequent time. Initial conditions had each species perfectly adapted to each of the alternate patch types as in the patch occupancy model and all patches filled with the species pre-adapted to the local environment. Disturbances were imposed randomly on patches, wiping out the local community, after which the empty patch could be recolonized. After colonization, a new locally adapted population could be established from pre-adapted immigrants and/or evolve from less-adapted ones, dependent on the relative speed of dispersal versus evolution.

Confirming the insights from the patch occupancy model, we found that regional neutrality involving all three of the characteristics we identified in the analytical model could evolve under a range of more realistic conditions (Figure 3–6) even though the simulations included numerous complications including explicit spatial structure, detailed genetic mechanisms, and stochastic effects of demography, dispersal and disturbance. We found that regional neutrality was more prevalent when dispersal was low (in our case when there were fewer than 100 individuals dispersing per generation [Figure 5]). It also takes longer to establish neutrality with lower extinction rates and higher dispersal rates. A difference between our simulations and the patch occupancy model is that our simulations could also generate trait convergence within patches that lead to local neutrality (as modeled by e.g. ter Horst et al. 2010), but we found that local cooccurence of species with identical trait values was rare and certainly much less common than evolutionary niche priority effects (Figure 5).

**Figure 3:**
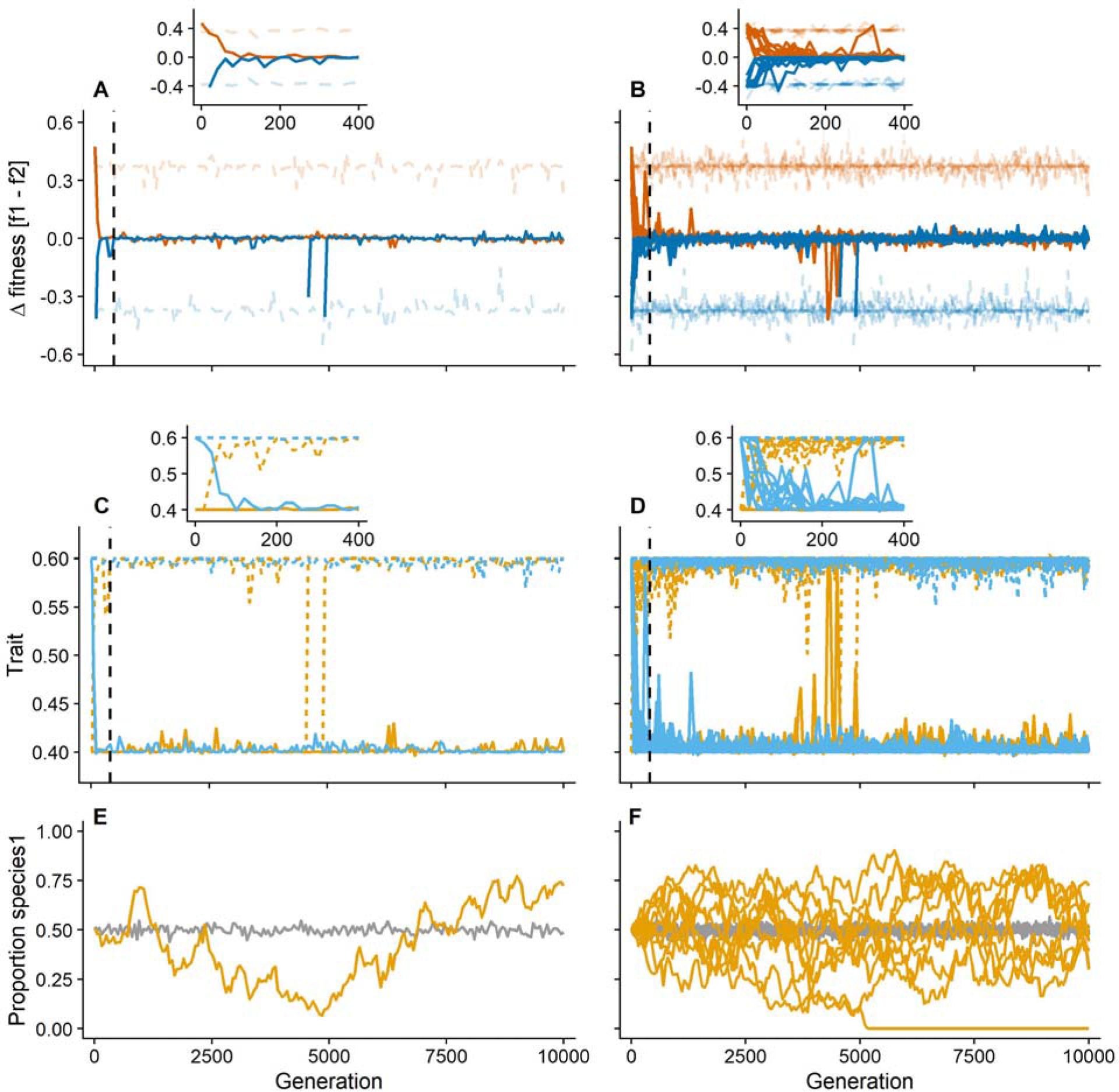
Temporal dynamics of evolution leading to regional neutrality. Results for sexual species with dispersal *Nm* = 10 individuals per generation and patch extinction rate *P*_*ext*_ = 0.01 per generation. Optimal trait values in each habitat are 0.4 and 0.6 in patch type A (yellow) and patch type B (blue) habitats respectively. **Left** (A, C, E) results for a single run, **Right** (B, D, F) results for ten replicate simulations. Small panes in A-D show a detailed view of the first 400 generations (indicated by vertical dashed line). **A,B**) Fitness difference between species 1 and species 2 (lighter dashed lines are control simulations without evolution) within each patch type (red in patch type 1, blue in patch type 2). Here evolution has led to mean fitness differences near zero within each patch type within the first 500 generations (inset). Note that there are several points in time in some of the simulations that show strong deviations from the general pattern. This occurs when due to drift and/or dispersal only few individuals of a species are present in one of the patch types and these are maladapted to that patch type. Often the species can reestablished itself more generally in that patch type in subsequent generations after this unique random event. **C,D**) Mean genotypic value for each species (orange = species 1, light blue = species 2) in each patch type (solid = patch type A, dashed = patch type B) through time. Here evolution has led to strong bimodal distributions in both species that link genotype to patch type within the first 500 generations (inset). **E,F**) Proportion of species 1 in the metacommunity through time (grey lines are control simulations without evolution). Evolution results in neutrality between species at the metacommunity level, which enhances drift in relative abundance of species in the metacommunity.

**Figure 4:**
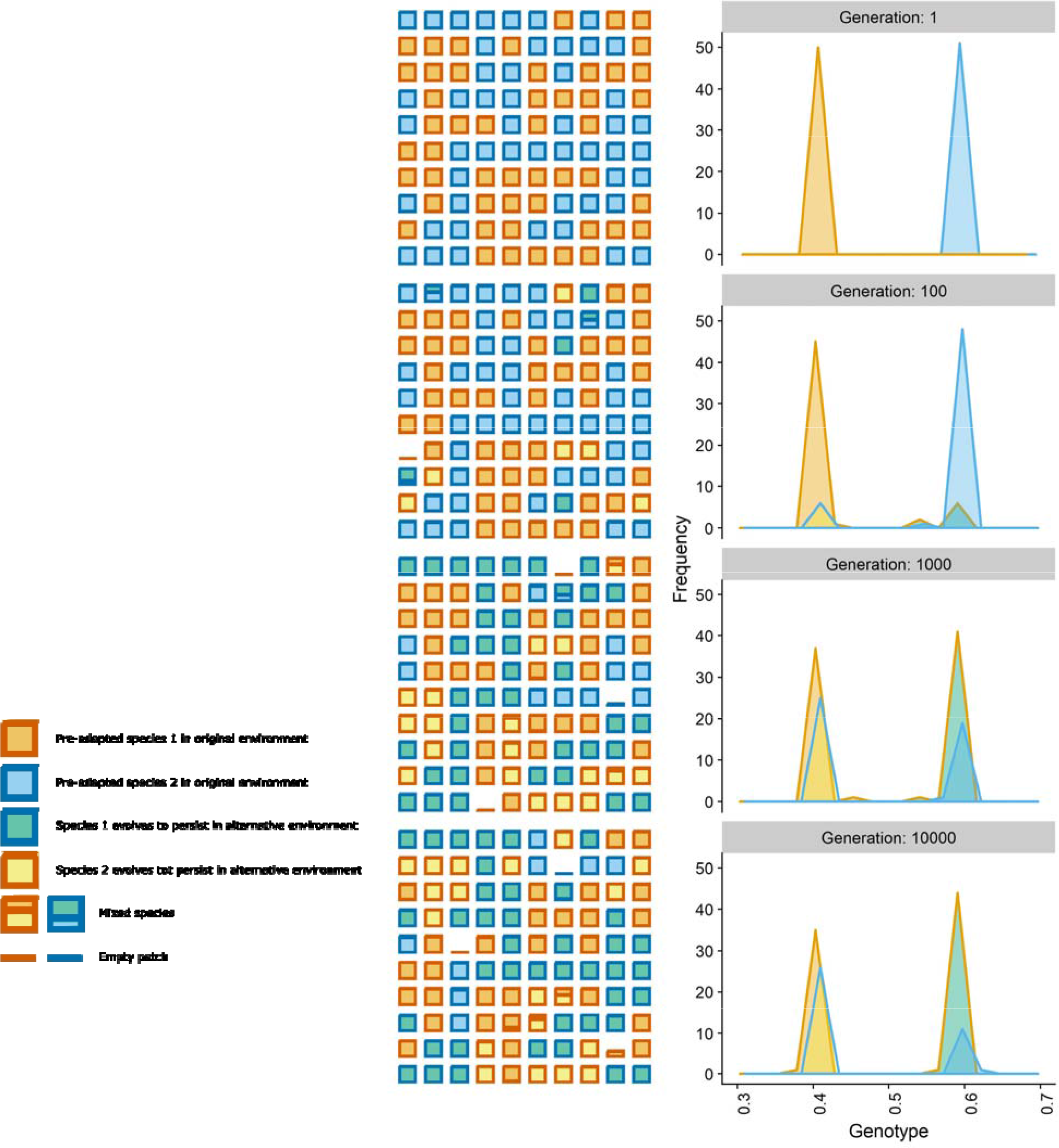
**Left**: patch occupancy of both species through time for a single run with sexual reproduction. Each square represents a habitat patch in the 10 by 10 grid. Edge color represents the patch type (red = patch type A, blue = patch type B). Fill color represents species × niche adaptation. In Generation 1 each species dominates in the patch type to which it is pre-adapted. With time, each species can dominate in either patch type due to evolution into the other niche. **Right**: Genotype distribution in the entire metacommunity of both species through time for a single run with sexual reproduction. Edge color represents species identity. Fill color represents species × niche adaptation. In generation 1 each species is pre-adapted to one of the patch types (trait values of 0.4 or 0.6). With time species evolve into each other’s niche, resulting in evolutionary convergence of similar bimodal niche trait distributions. Results for dispersal *Nm* = 10 individuals per generation and patch extinction rate *P*_*ext*_ = 0.01 per generation.

**Figure 5:**
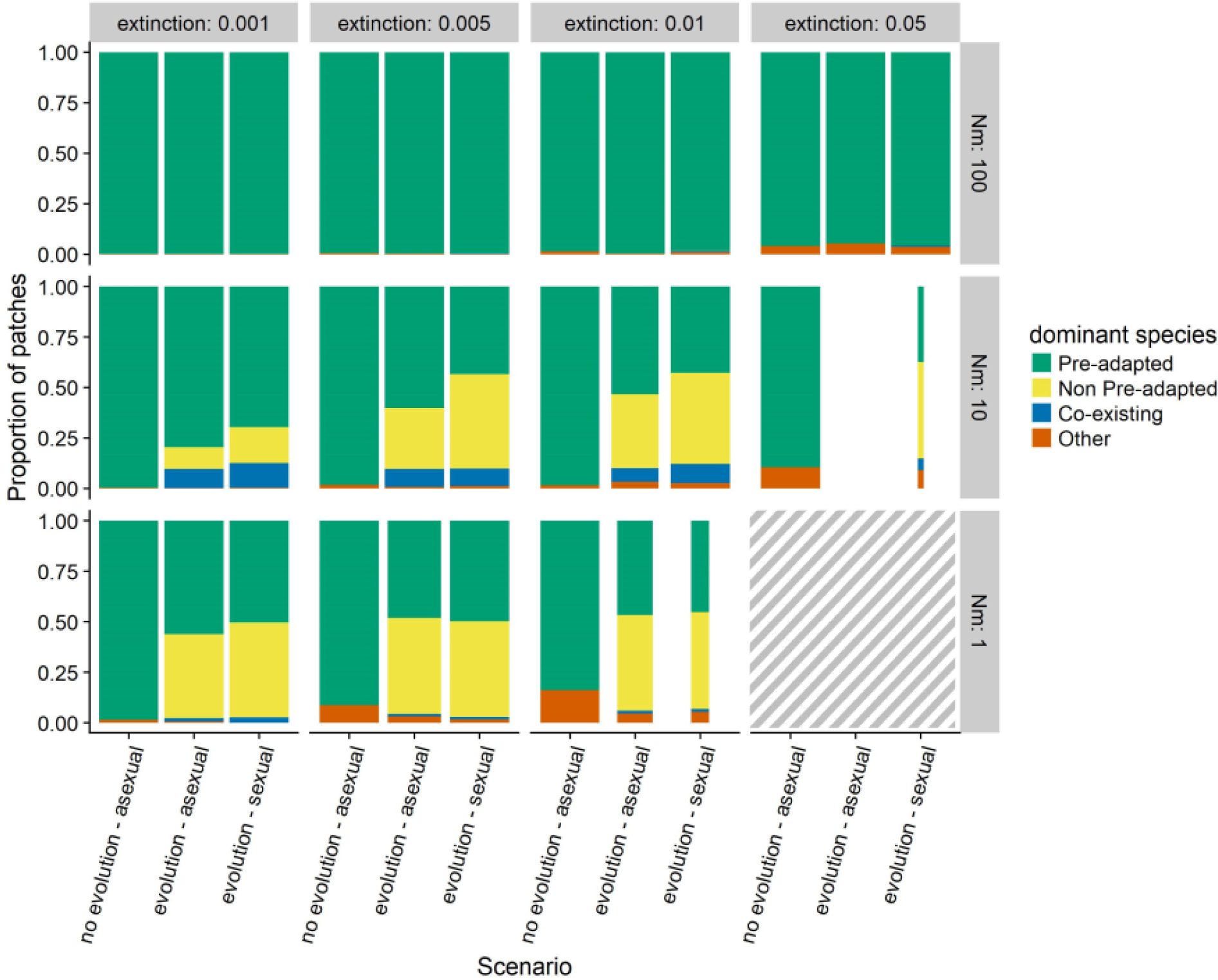
Patterns of patch dominance at generation 5000 by pre-adapted or non pre-adapted species for different levels of patch extinction and dispersal. Results are averaged over 10 replicate runs. Color codes do not represent species or patch type, but indicate whether the dominant species in a patch was pre-adapted to the patch type at the start of the simulations. Green shows patches occupied either by Species 1 with trait values corresponding to A patches in A type patches or by Species 2 with trait values corresponding to B patches in B type patches. These correspond to patches expected to be dominated by the specialist species for each patch type in the model without evolution. Yellow shows patches occupied by either Species 2 with trait values correspond to A patches in A-type patches or by Species 1 with trait values corresponding to B patches in B-type patches. These correspond to cases where species have strongly switched trait values due to evolution so they are dominant in the habitat in which they were initially subordinate. Such patches do not exist in the absence of evolution. Blue shows patches where both species co-exist and are both adapted to the local environment. Brown shows all other cases including where patches are occupied by populations of either or both species that are not well adapted to local conditions or patches that are empty. Width of the bars indicates the proportion of replicate runs (total of 10) where both species still co-exist in the metacommunity and none of the species has gone extinct due to drift after regional neutrality has established. Hatched area indicates parameter settings where both species go extinct.

Another important difference between our simulations and the patch occupancy model is that our simulations include stochastic processes that reveal the eventual role of ecological drift at the regional scale on the relative abundances of the two species (Figure 3E,F). This occurs in our simulations because of the finite size of the metacommunity (finite number of patches and consequent finite number of individuals) which was absent in the patch occupancy model that assumed an infinite number of patches. Given a finite metacommunity size, the relative occupancy (number of sites occupied) and abundances will eventually lead to the extinction of one or the other species even if this can take a very long time. The extent of drift will depend on the size of the metacommunity (patch number), extinction rate and dispersal (Figure A8 in appendix).

We next used a standard statistical tool (‘variation partitioning’ sensu Peres-Neto et al. 2006) used by ecologists to discern what mechanisms might underlie community patterns in nature. This tool partitions the variance in community composition to that explained by environmental and spatial variation, as well as variation common to the two (confounding of both effects) and an unexplained element largely attributable to stochasticity or unmeasured environmental components (absent in our model where we specify the environment). When environmental variation is randomly structured in space, variance explained by the environmental component is usually attributed to niche-based processes whereas the spatial and unexplained component are assumed to reflect a combination of dispersal limitation and drift. We applied this technique to our simulation data and found that evolved regional neutrality substantially reduced contributions from niche-based processes and inflated the spatial and residual variation (Figure 6). If we do a similar partitioning of trait variation (rather than community composition) we find that environments explain most of the variation in traits whereas spatial and residual effects are small. These results show that purely neutral (scale independent) ecological processes and local niche based evolution of regional neutrality cannot be separated by this method without also looking at trait variation.

**Figure 6:**
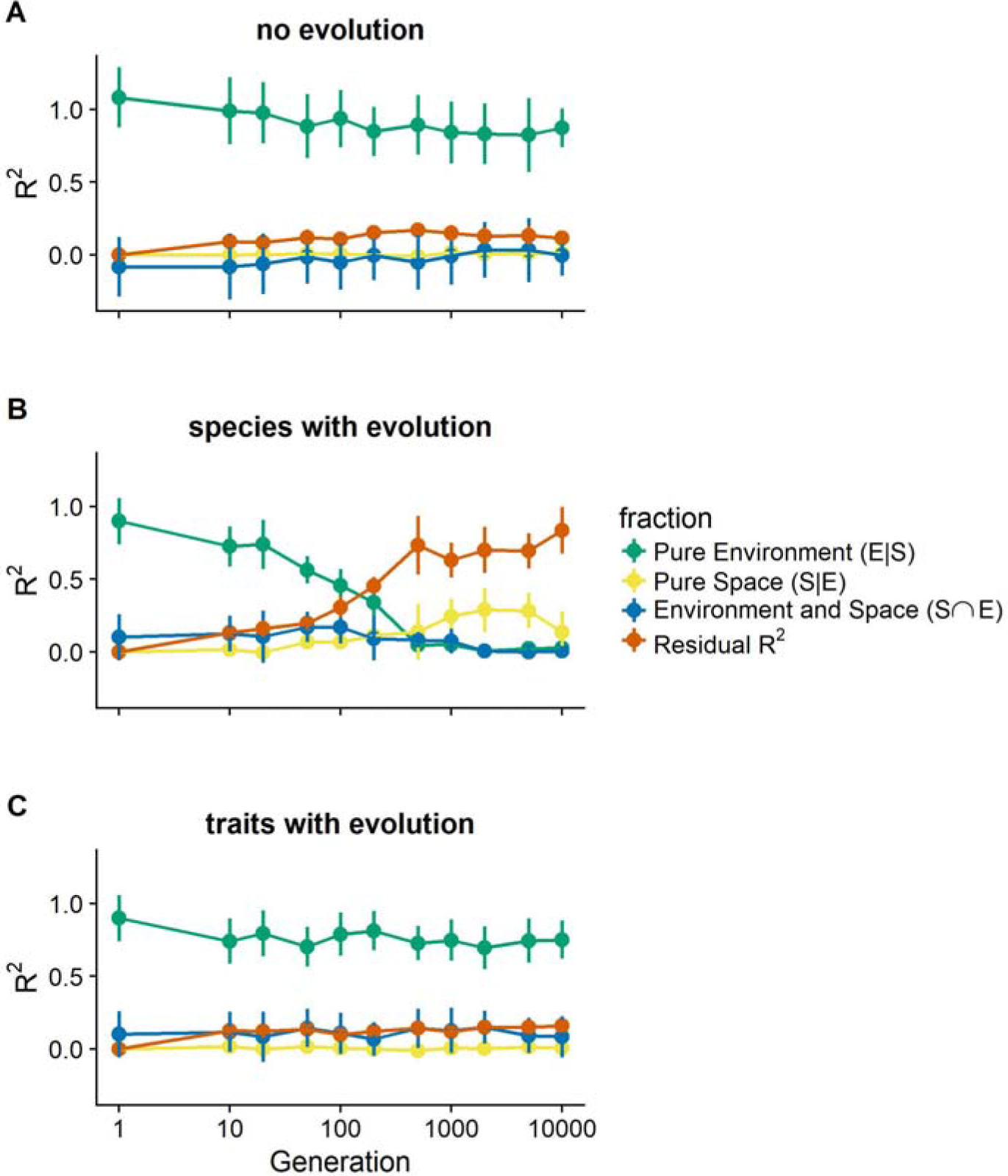
Variation partitioning components through time for species and for traits. Results for dispersal *Nm* = 10 individuals per generation and patch extinction rate *P*_*ext*_ = 0.01 per generation. **A**) results in the absence of evolution (controls) for species and for traits. **B**) results for species for dynamics with evolution. **C**) results for traits for dynamics with evolution. In the absence of evolution species and traits are equivalent. With evolution, the results for traits are very similar to control simulations but the pattern for species shows a very strong decline in effects of environment and a strong increase in residual and spatial effects by generation 1000.

Although there are quantitatively different results, we nevertheless find that regional neutrality, linked to local evolutionary priority effects as in the two species case described above, also occur in simulations with asexual reproduction (Figure A1), broader dispersal kernels (Figure A2), weaker selection (Figure A3), lower reproduction rates (Figure A4) and more (four) species (Figure A7). We also find that similar effects occur when patch distributions are uneven (i.e deviate from the 1:1 ratio we used here) although they increasingly lead to the extinction of one of the species early in the simulations (Figures A5 and A6).

## Discussion

Our results reveal that niche-driven, localized adaptation in a heterogeneous landscape can lead to patterns of regional neutral equivalence of species. If so, the dichotomous tension between neutrality and niche partitioning suggested by purely ecological models (Gravel et al. 2006, Leibold and McPeek 2006, Adler et al. 2007) may be misleading because the two effects can be scale dependent. At the scale of the local patch, the dynamics determining the relative abundances of species in our models are strongly niche-driven while phenotypic variation and fitness profiles reveal ecological equivalence among species at the regional level. Thus, patterns that appear neutral at the regional scale might result from ongoing and strongly niche-driven dynamics in different patches mediated by both evolutionary (genetic adaptation to the local habitat type) and ecological (changes in the relative abundance of species) processes.

Regional neutrality makes novel predictions about the structure and dynamics of ecological communities in heterogeneous landscapes. Under regional neutrality we predict adaptive divergence among populations within species at the local scale and geographic convergence of fitness-related traits and subsequent neutral drift of species toward a single dominant one at the regional scale. With uneven environmental distributions, regional niches will reflect environmental distributions and drift will be biased toward the species initially adapted to the more prevalent habitat (Appendices Figure A6 and A7). Regional neutrality requires that selection is not so severe as to prevent establishment of maladapted populations, and repeated disturbances that provide the template on which these populations can adapt and expand the regional niche.

One of the more important predictions to emerge from the regional neutrality hypothesis is that taxa will show little environmental tracking and potentially show spatial patterning, whereas local variation in trait values (independent of taxa) will show strong environmental tracking and less spatial patterning. This contrasts with more conventional predictions from purely ecological species sorting (no local adaptation) where both taxa and traits should show environmental tracking, and from conventional predictions based on neutrality involving purely ecological processes where neither species nor traits will show environmental tracking and both show similar spatial patterning. Of course, this will involve measuring traits at the population rather than species level and will depend on there being a close connection between the traits and the fitness response to environment. Unfortunately we know of no studies that have examined such patterns in sufficient detail to be illustrative of this point.

Our model is not the first to claim that local adaptive evolution can lead to neutrality but it differs in important ways from these previous efforts. Scheffer & van Nes (2006) showed that competitive interactions along an environmental gradient can lead both to niche partitioning as well as to the emergence of subgroups of ecologically equivalent species. This is a mechanism leading to evolution of neutrality at the local patch scale. In our simulations, the emergence of neutrality at the local scale also occurs in a generally similar way but it is relatively rare. Instead we find that the interaction of local evolution, dispersal and species sorting lead to a much stronger pattern of ecological equivalence of species at the regional scale and to much more frequent niche specialization at the local scale. The difference is that we incorporate recurrent local disturbances that iterate the process of local adaptive niche evolution and this allows species to eventually colonize alternative habitat types, adapt, and expand the species’ overall niche in ways that cannot happen in environmentally fixed landscapes (no disturbances or local environmental fluctuations). Hubbell (2006) finds that spatially autocorrelated environments and random initiation of species can lead to the evolution of habitat generalists with locally distinct niche traits, but his work did not reveal if these conditions are sufficient to obtain regional neutrality in the model (e.g., competitive equivalence and eventual neutral drift). His work also suggests that these results would be most likely in highly diverse communities, while we show that this outcome is possible even for pairwise interactions. Our approach also differs somewhat from that of Hubbell (2006) by having discrete patches rather than continuous environmental gradients. We imagine that adaptive dynamics in such cases might differ somewhat but this is difficult to evaluate with the patch-like structure of our model.

Both of our models are a highly simplified caricatures of the natural world. A variety of features could decrease the degree to which regional neutrality occurs in natural settings. For example, we assume that both species can evolve the optimal trait and at similar rates. Lower additive genetic variation and other genetic constraints could limit one or both species in this regard or even result in regional exclusion. However, many analyses suggest that ecologically important traits can evolve rapidly, and more closely related species, such as those that are likely to compete for the same resources, might be more likely to have similar evolutionary rates and capacities.

Local adaptive evolution might often contribute to cryptic eco-evolutionary dynamics that alter the regional mechanisms that determine biological diversity and resistance to disturbance. The evolution of regionally neutral species means that drift could reduce diversity over long periods as stochastic events lead one species to dominate and the other to become extinct. Yet, regionally neutral species would also provide a level of redundancy during perturbations. For instance, if these species contribute to ecosystem function, then losing one species would still maintain this function across all habitat types. Moreover, as habitats change through natural or anthropogenic means, local adaptive evolution could keep pace allowing each species’ regional niche to expand given sufficient additive genetic variation. Fully understanding the contribution of evolution to biodiversity patterns will require a more integrated biology that synthesizes community ecology and evolutionary biology across a range of temporal and spatial scales.

## Methods

### Patch occupancy model

We keep track of occupancy *p* of two patch types (*env*=A,B) by two species (*sp*=1,2) with two optimal patch type phenotypes (*opt*=A,B), denoted by *p*_*sP,opt,env*_. Empty patches of type *env* are given by *p*_0,*env*_ = *h*_*env*_ − Σ_*sp*_ Σ_*opt*_ *p*_*sp,opt,env*_ with *h*_*env*_ the proportion of the landscape with patch type *env*. Empty patches are invasible by all combinations of species *sp* and phenotype *opt*. Occupied patches are invasible if phenotype *opt* of the resident does not match patch type *env* and phenotype *opt* of the invader does. When invaded, the entire population is instantly replaced by the invader (no explicit population dynamics). The patch dynamics are given by:

**Species 1:**

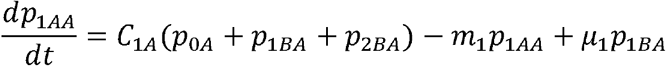

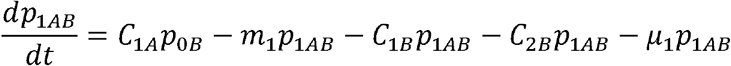

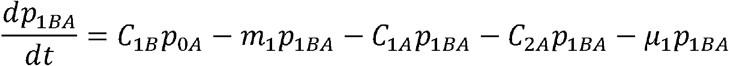

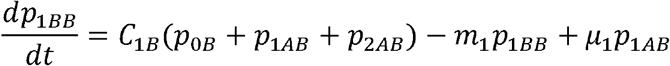
**Species 2:**

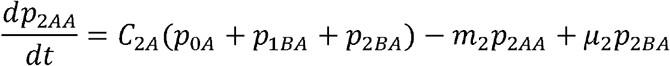

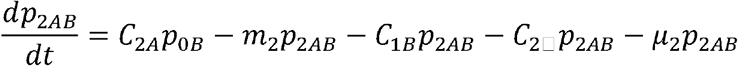

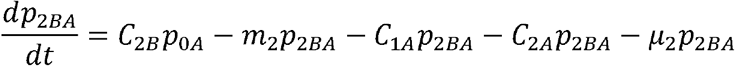

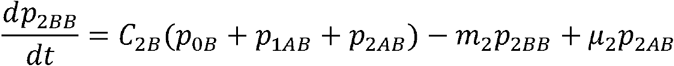

where *C*_*sp,opt*_ = Σ_*env*_ *c*_*sp,opt,env*_ *p*_*sp,opt,env*_ denotes the overall propagule production of species *sp*, phenotype *opt*.

To assume regional neutrality, let *m*_1_ = *m*_2_ = *m*, *c*_1,*opt,env*_ = *c*_2,*opt,env*_ = *c*_*opt,env*_ and *μ*_1_ = *μ*_2_ = *μ*.

To assume symmetry between patch types, let *h*_*A*_ = *h*_*B*_ = 1/2 and

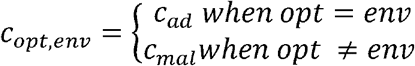

We initialize simulations with species 1 adapted to patch A and species 2 adapted to patch B (*p*_1*BA*_(0) = *p*_1*BB*_(0) = 0,*p*_2*AA*_(0) = *p*_2*AB*_(0) = 0,*p*_1*AA*_(0) = *p*_1*AB*_(0) = *p*_2*BA*_(0) = *p*_2*BB*_(0) = 0.01,*p*_*0A*_(0) = *p*_*0B*_(0) = 0.48). The dynamics of the model are solved numerically using NDSolve in Mathematica.

### Spatially explicit IBM

We create a 10 × 10 square grid of interconnected patches inhabited by species competing for available resources. There are 2 different environments that are randomly distributed among the patches (no correlation between environment and space), with equal representation of each environment. For each of the two environments there is a pre-adapted species present. Species have discrete reproduction, overlapping generations and newborn dispersal and can be either sexual or asexual (with the restriction that both species have the same reproductive mode). Patches are linked by dispersal of individuals from all species present.

Within each patch, individuals inhabit microsites with environment *e* equal the mean patch environment plus stochastic Gaussian variance 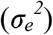. Each individual has a phenotype *Z*_*i*_ that determines its fitness and that consists of a genetic component and some random non-genetic contribution according to a Gaussian distribution (mean = 0, variance 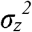) determined at birth. The genotypic value ranges from zero to one and is calculated as the mean of *L* = 20 bi-allelic additive genes (value 0 or 1). Fitness of the individuals is calculated as

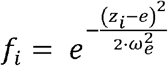

where *e* is the environment in the microsite inhabited by individual *i* and *ω*_*e*_ is the width of stabilizing selection. Larger values of *ω*_*e*_ result in weaker environmental selection and smaller values of *ω*_*e*_ result in stronger environmental selection. The probability of survival *Si* of an individual at each time step is calculated according to

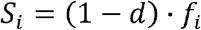

where *d* = 0.1 is the baseline mortality. Following survival, the number of offspring individuals *I*_*p*_ to be assigned to each patch *p* is calculated as

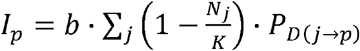

where *b* is the per capita birth rate (*b* = 2), *N*_*j*_ is the number of individuals (pooled over species) in patch *j*, *K*is the carrying capacity and *P*_*D(i→p)*_ is the probability of offspring dispersal from patch *j* to patch *p* (including no dispersal when *j* = *p*). If *N*_*j*_ is larger than *K*, 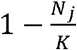 is set to zero. If patch *j* and *p* are the same *P*_*D(j→p)*_ equals (1 − *m*), where *m* is the dispersal rate. If patch *j* and *p* are different patches, *P*_*D(j→p)*_ follows an exponential decay model and is calculated in two steps. First the exponential decay is calculated in function of the distance between patch *j* and *p* as

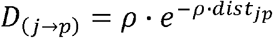

where *ρ* is a scaling factor determining the strength of dispersal limitation and *dist*_*jp*_ is the Euclidean distance of grid coordinates between patch *j* and *p*. The larger *ρ*, the narrower the dispersal kernel. Next the dispersal rate between patch *j* and *p* is rescaled as

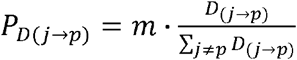

such that the total dispersal rate from all other patches to patch *p* equals *m*.

Based on the probabilities, 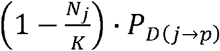, mothers are randomly drawn with replacement for each of the *I*_*p*_ offspring. In case of sexual reproduction, a matching father is randomly drawn with replacement from the same patch and the same species. Individuals are hermaphrodite and self-fertilization is allowed, such that there is no demographic cost of sex.

Asexually produced offspring inherit the entire genome from their single parent without recombination. Sexually produced offspring inherit from each parent one randomly selected gene for each of the 10 diploid loci. Both for asexual and sexual reproduction mutation occurs with probability *μ* at each gene.

At *t* = 0, all patches are filled with the species pre-adapted to the local environment. During the simulations there is a given probability *P*_*ext*_ of patch extinction. If a patch goes extinct, it needs to be recolonized from the other patches in the metacommunity. We ran 10 replicate simulations for a range of extinction rates (*P*_*ext*_) and dispersal rates. For fully adapted individuals, generation time equals 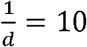 time steps. Simulations are run for 100000 time steps or 10000 generations. Dispersal is expressed as the absolute number of offspring dispersing per generation among patches 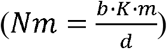.

## Supplemental Information

Our results are robust to variation in most of the parameters we investigated. The figures below show results for simulations that differ from our baseline scenario as follows: In each case the figures should be compared with Figures 3–6 in the main text.

**Figure A1:**
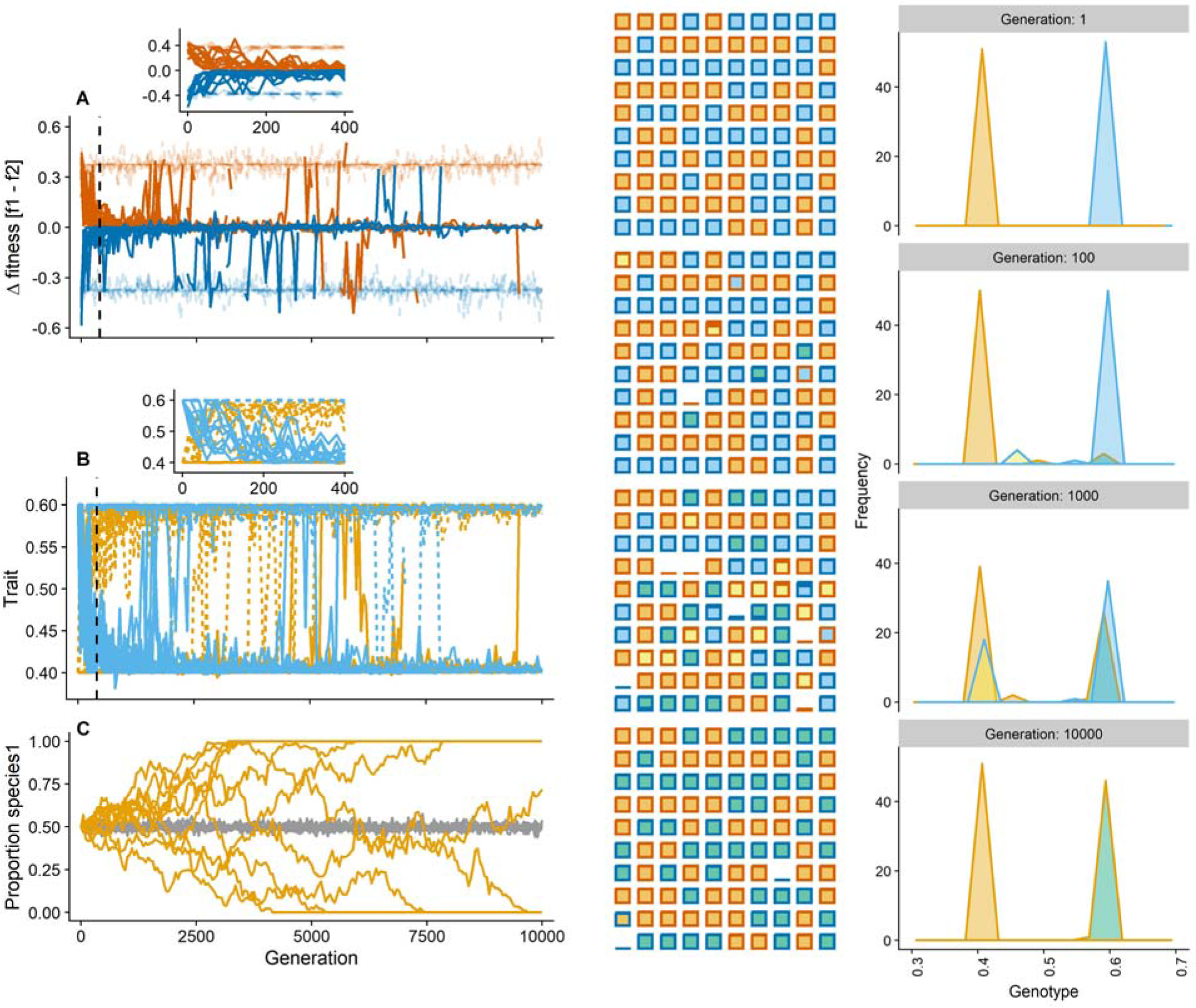
Simulations with asexual reproduction instead of sexual. See Figures 3 and 4 for legend details. The time to reach regional neutrality was longer (slower evolution) and the amount of drift was higher than compared to sexual reproduction (Compare with Figures 3 and 4).

**Figure A2:**
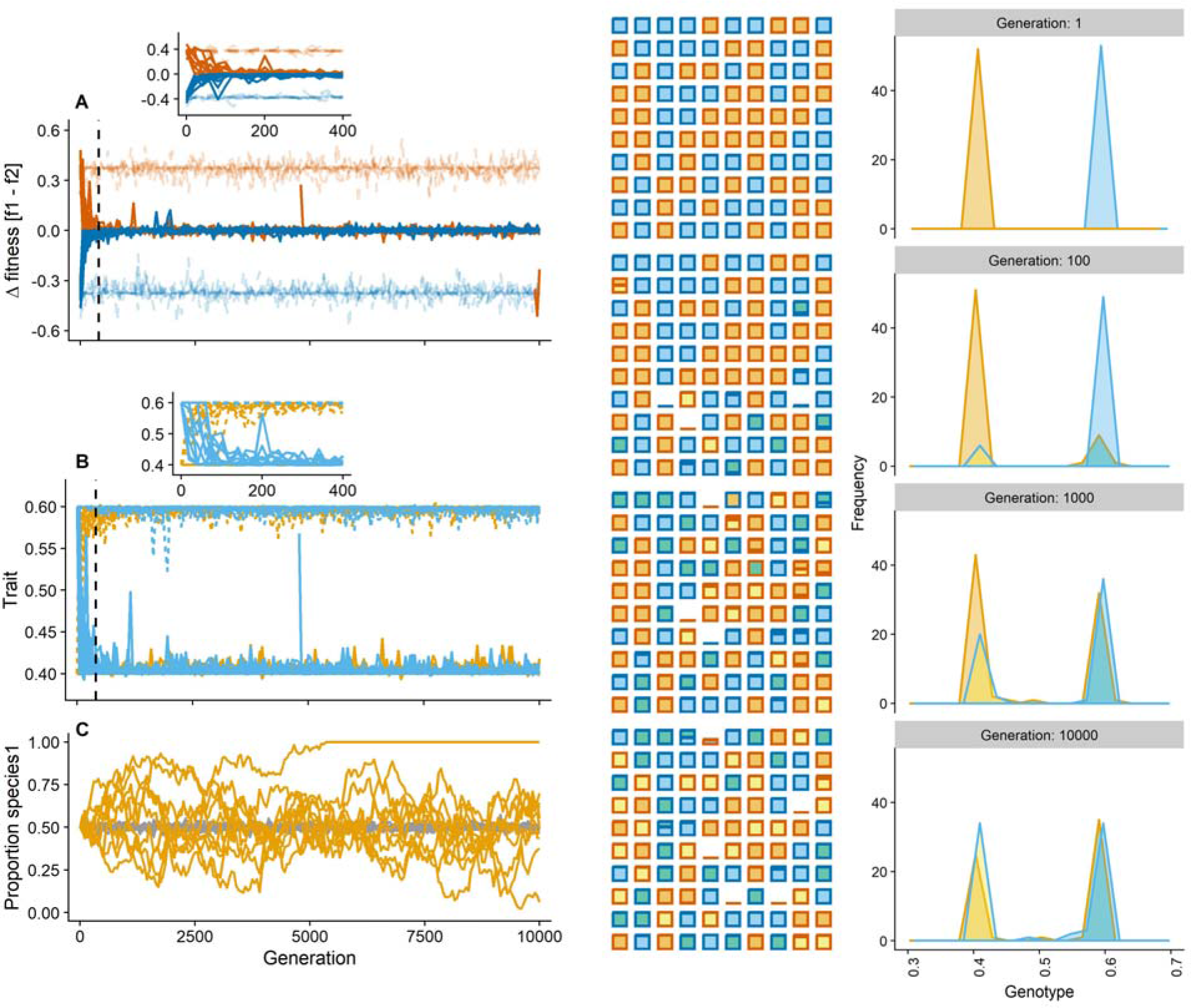
Simulations with broader dispersal kernel (*ρ* = 0.2). See Figures 3 and 4 for legend details. There were no noticeable effects compared with baseline conditions (Compare with Figures 3 and 4).

**Figure A3:**
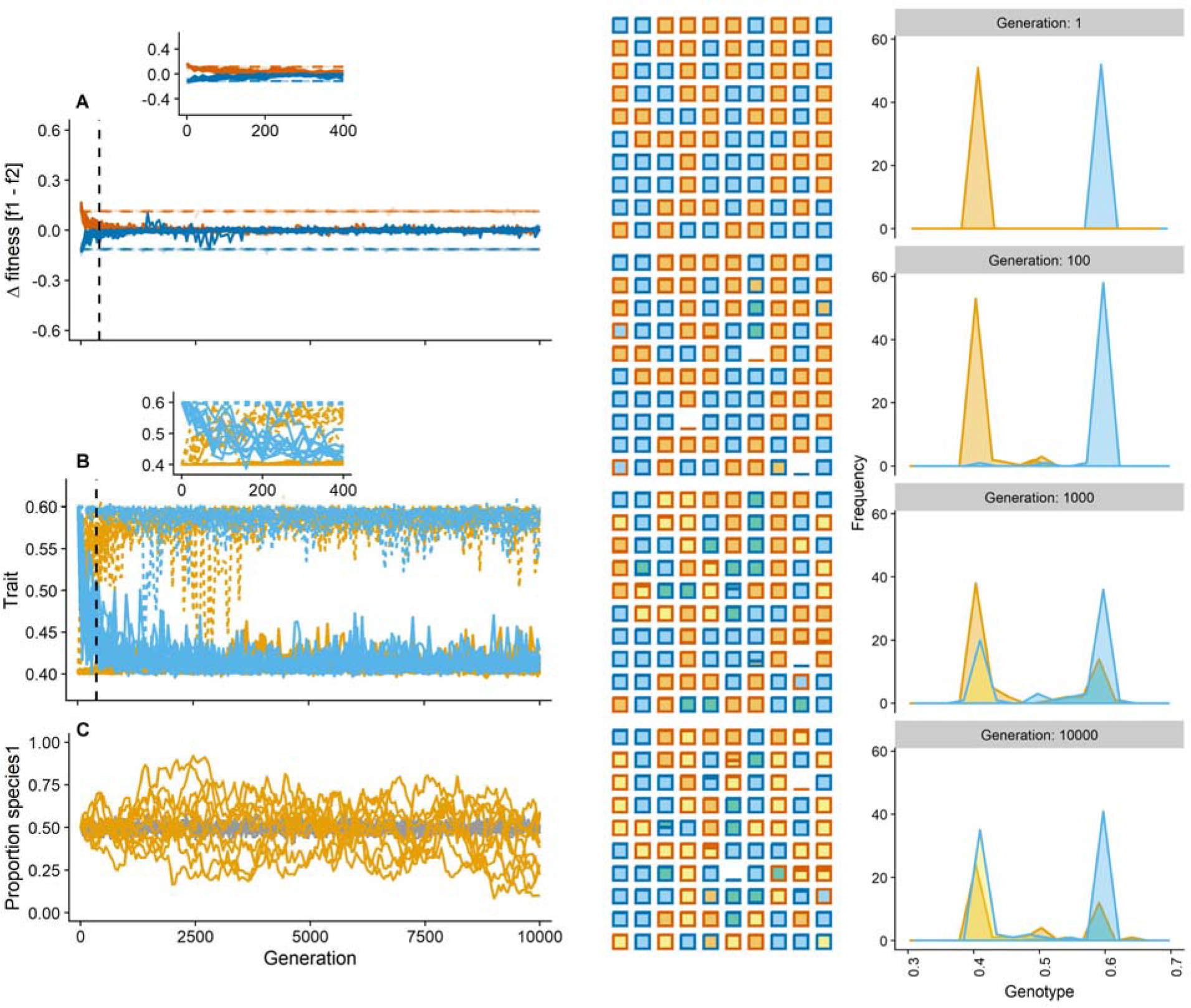
Simulations with weaker selection difference between habitats: (*ω*_*e*_ = 0.4). See Figures 3 and 4 for legend details. Evolution is slower due to lower fitness differences between environments (non-adapted species have less disadvantage) but colonization and regional dynamics are similar to baseline conditions (Compare with Figures 3 and 4).

**Figure A4:**
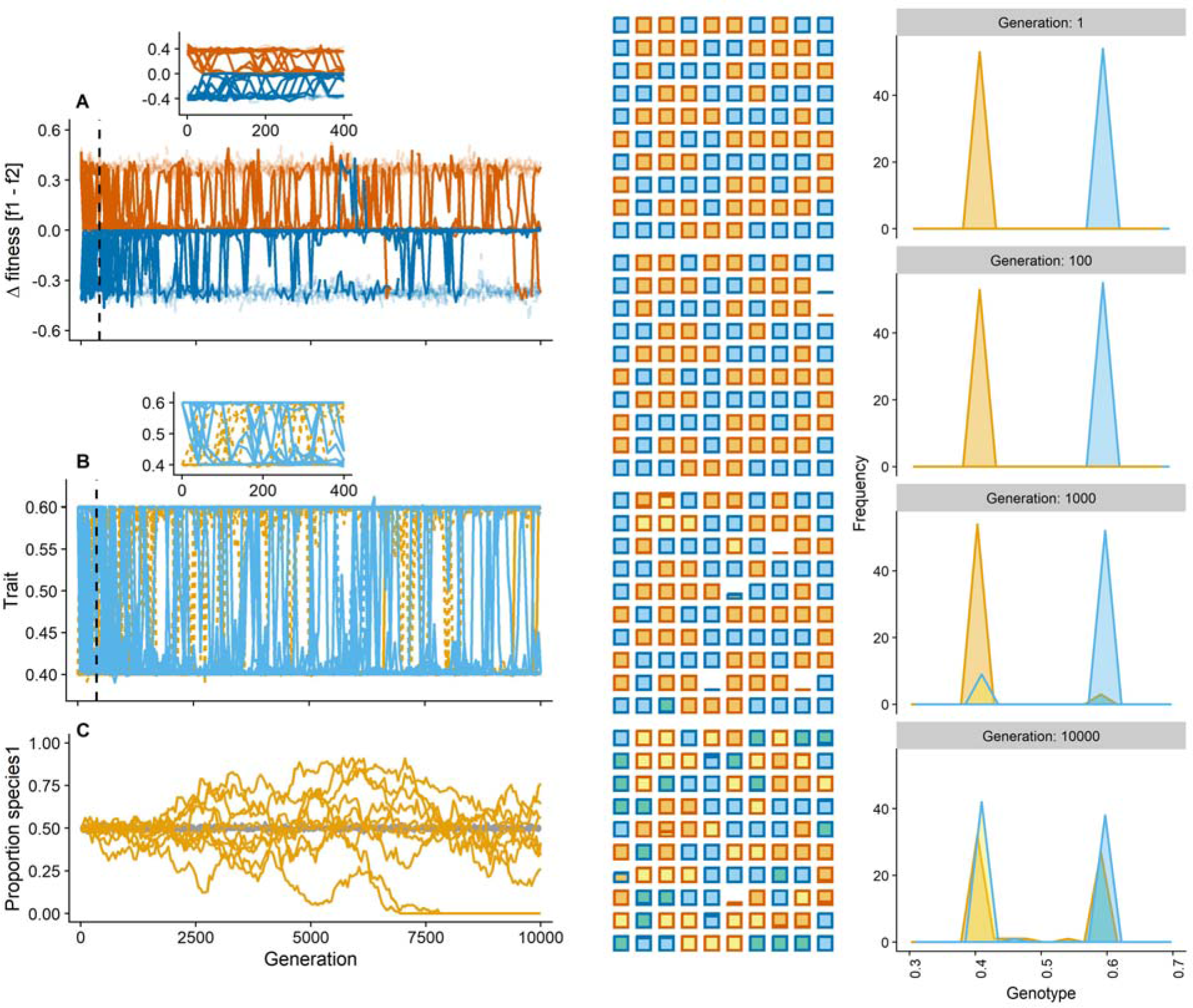
Simulations with lower reproduction rate: (*b* = 1). See Figures 3 and 4 for legend details. The degree of regional neutrality was slower and there were more erratic (but recoverable) extinctions of species from one of both environments. (Compare with Figures 3 and 4).

**Figure A5:**
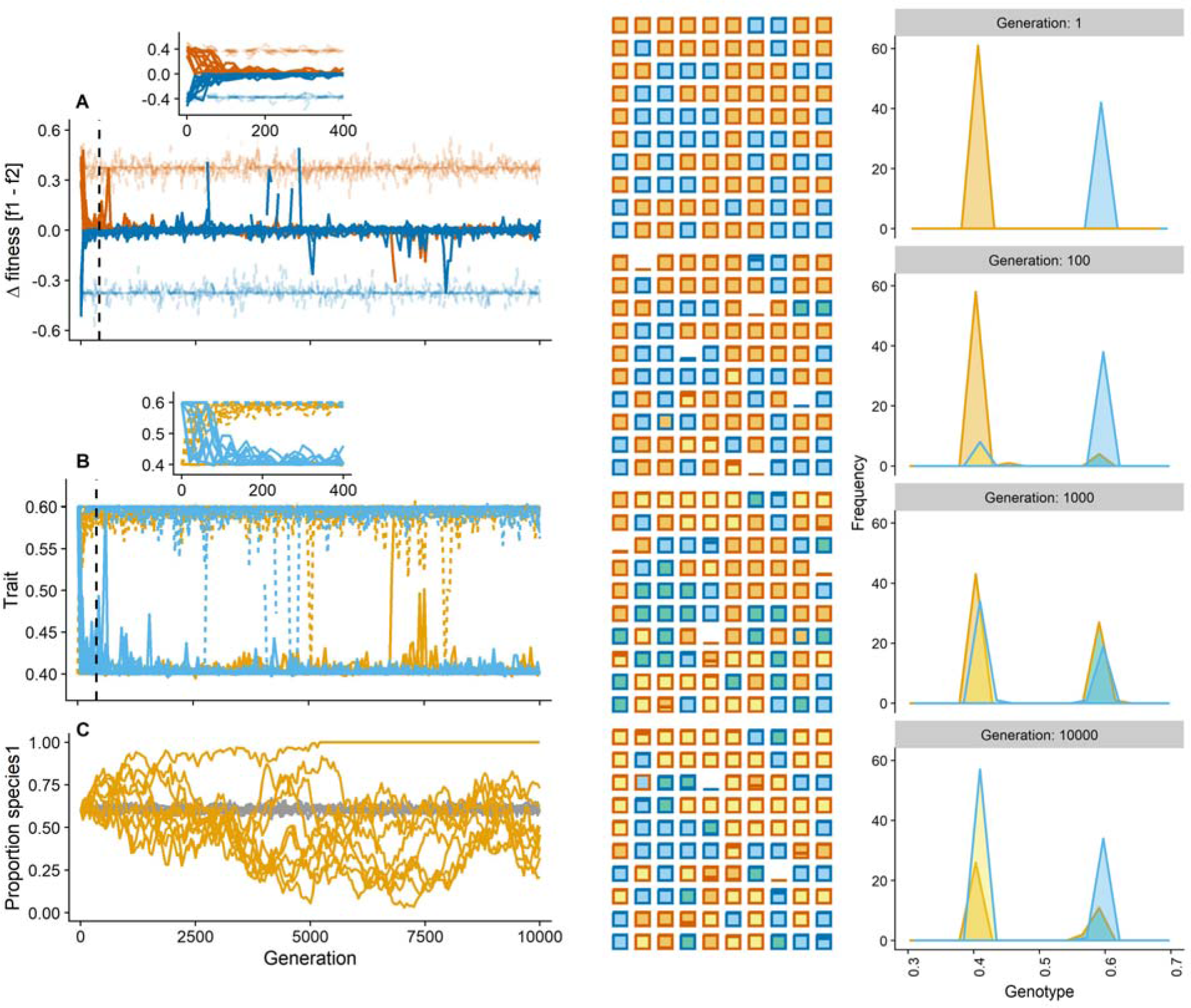
Simulations with asymmetric patch type distribution: 60% of patch type A, 40% type B. See Figures 3 and 4 for legend details. There was a tendency for species 2 to go extinct (one replicate) early in the simulations (Compare with Figures 3 and 4).

**Figure A6:**
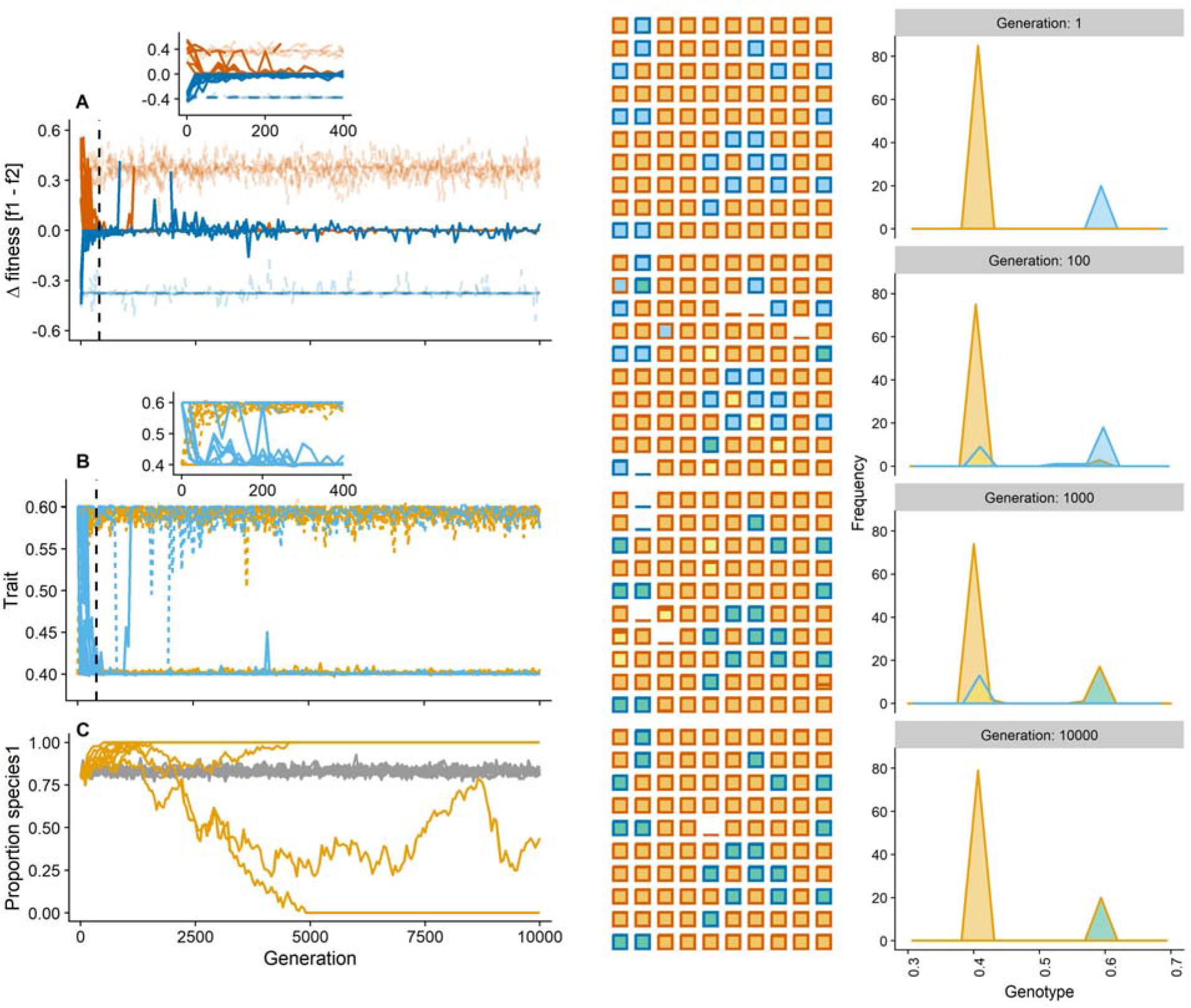
Simulations with stronger asymmetric patch type distribution: 80% of patch type A, 20% type B. See Figures 3 and 4 for legend details. There was a strong tendency for species 2 to go extinct (8 replicates) than baseline conditions (Compare with Figures 3 and 4).

**Figure A7:**
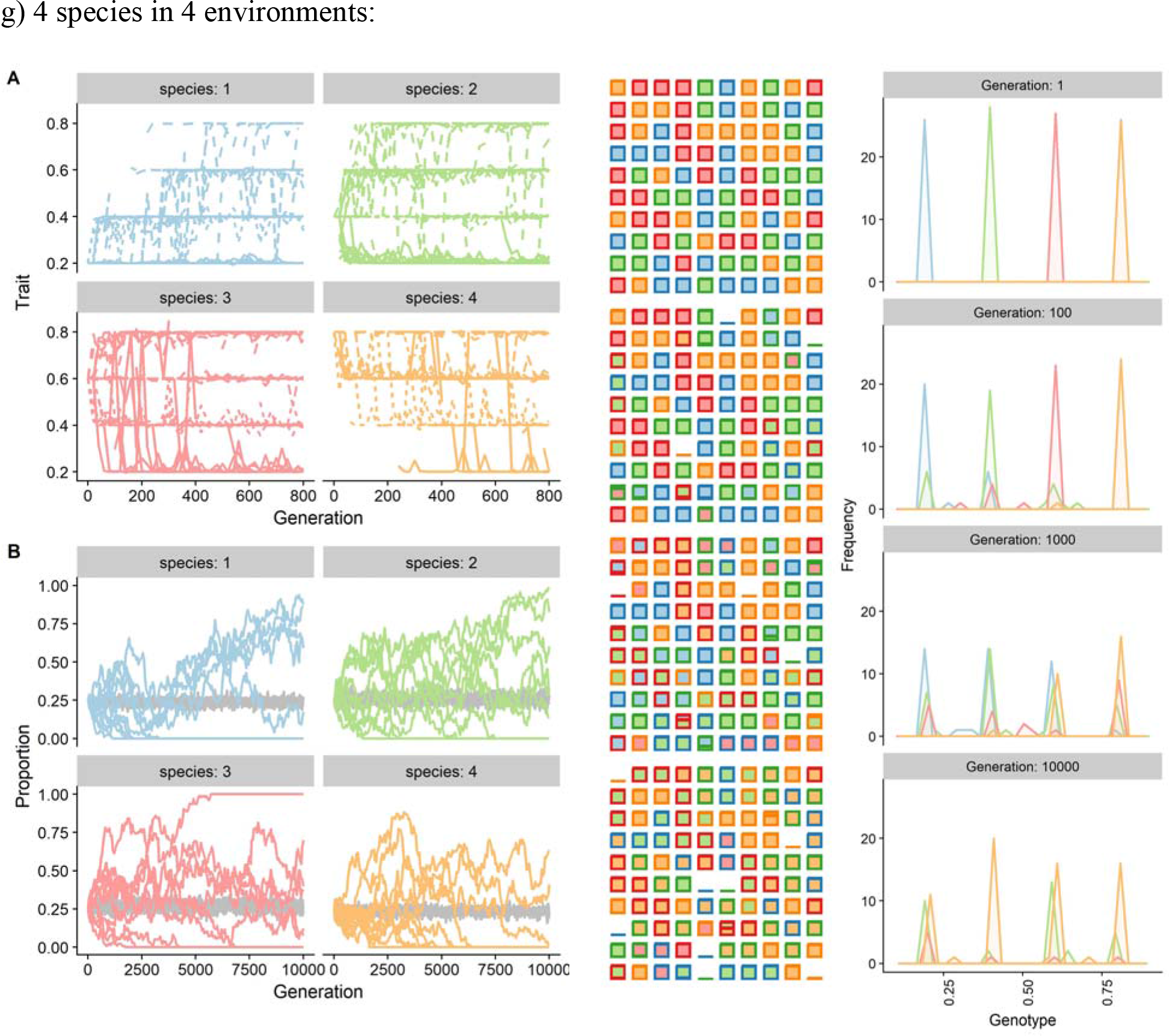
Simulations with 4 species arrayed and 4 patch types along a single trait gradient.
See Figures 3 and 4 for legend details. The leftmost figure (comparable to Figure 3) is structured slightly different and lacks the panel for fitness difference between the species. In the middle figure (patch occupation), fill color represents species identity and not species x niche adaptation. In the rightmost figure (genotype frequencies), both edge and fill color represent species identity (and not species x niche adaptation). All four species eventually evolved to specialize on every patch type (at least to some degree). The dynamics were much slower however and were still changing at the end of the simulation. There was also more drift and extinctions than baseline. (Compare with Figures 3 and 4).

**Figure A8:**
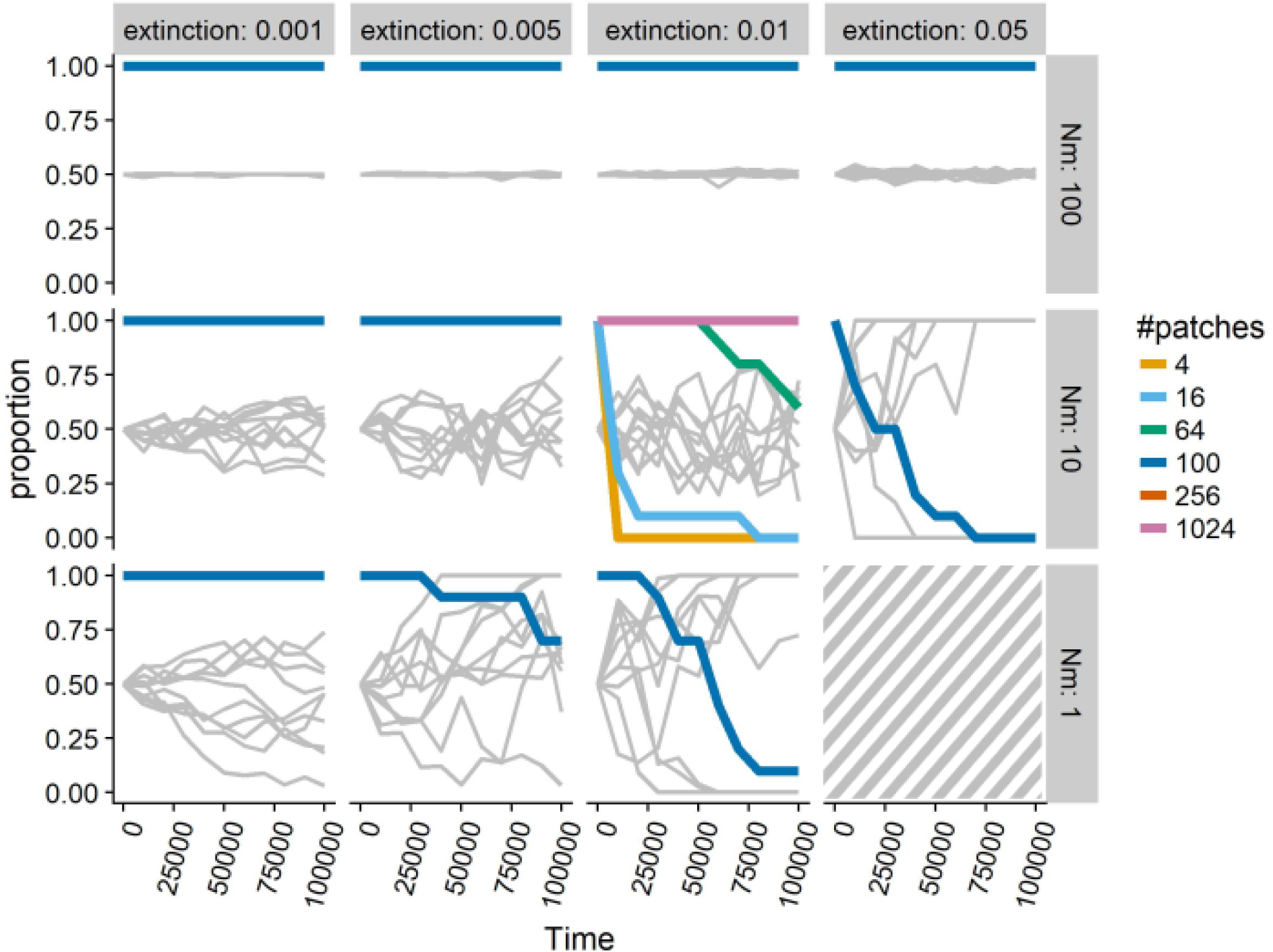
Interactive effects of extinction rate, dispersal and size of the metacommunity (patch number) on the extent of drift. Grey lines represent the proportion of species 1 in the metacommunity for 10 replicate runs under basline conditions (f# patches = 100). Colored lines represent the proportion of runs with >1 species in the metacommunity (each run starts with 2 species at proportion 0.5) for different metacommunity sizes. Different metacommunity size (patch number) is only modeled for extinction rate = 0.01 and dispersal = 10. Hatched area indicates parameter settings where both species go extinct. Drift increases with lower dispersal, higher extinction rates and smaller metacommunity sizes.

